# Frequency but not phase specific modulation of binocular rivalry with transcranial alternating current stimulation

**DOI:** 10.1101/690800

**Authors:** Jorge Delgado, Guillaume Riesen, Vladimir Y. Vildavski, Anthony M. Norcia

**Affiliations:** Biology Undergraduate Program, Stanford University; Neurosciences Graduate Program, Stanford University; Department of Psychology, Stanford University

**Keywords:** Transcranial Electrical Stimulation, Transcranial Alternating Current Stimulation, visual perception, binocular rivalry

## Abstract

Recent transcranial alternating current stimulation (tACS) literature suggests that tACS effects can in principle be both frequency and phase specific. In a series of three experiments using 69 participants used binocular rivalry percepts as a read-out for the effects of phase-synchronized tACS stimulation. To test for phase specificity, with frequency the same in each eye, we visually stimulated each eye with 3Hz, with stimuli in each eye presented in temporal in antiphase. The frequency-specific paradigm visually stimulated the right eye with 3Hz, and the left eye with 5Hz. Each experiment was accompanied by 3Hz tACS, whose phase with respect to the visual stimulus was varied by 0°, 90°, 180°, or 270° in relation to the right eye’s stimulus. A baseline no-tACS block preceded the stimulation blocks and two more followed, immediately and ten minutes after. Individual blocks lasted 4 minutes. Additionally, a no-tACS control experiment identical to the 3 Hz anti-phase visual stimuli setup was conducted, keeping all parameters the same but eliminating tACS. During stimulation, the 3 Hz anti-phase visual stimuli setup slowed the rate of rivalry in both eyes. Conversely, the 3Hz-right, 5Hz-left setup slowed the right (targeted) eye significantly while leaving the left (unstimulated) eye unchanged. In both experiments, durations returned to baseline after 10 minutes. Our results are consistent with the frequency-specific model of tACS, and with the Leveltian hypothesis that stimulation weakens the stimulated eye, as the right eye got weaker when it was directly targeted, and both eyes got weaker when targeted in antiphase. tACS does not appear to preferentially modulating percept durations in one phase more than in another.

## INTRODUCTION

Neural activity can be modulated by electric fields generated by electrodes placed on the scalp – a technique referred to as transcranial electrical stimulation (tES). Three commonly used tES methods differ in the temporal profile of the applied current, *e.g.* direct current (transcranial direct current stimulation; tDCS), alternating current (transcranial alternating current stimulation; tACS), or random noise currents (transcranial random noise stimulation; tRNS). In comparison to TDCS or TRNS, tACS is a particularly effective method to address possible modes of action as it can be delivered at frequencies that are designed to interact both with natural brain-wave rhythms (Antal and Paulus, 2013; Helfrich et al., 2014) as well as responses to externally applied periodic sensory stimuli (Neuling et al., 2012).

Prior work on the mechanism of action of tACS has used invasive recordings in animal models and has suggested several mechanisms by which the imposed tACS field could affect with ongoing neural activity, either instantaneously (Liu et al., 2018) or as a (partially) sustained after-effect (Stagg et al., 2018). One of the proposed instantaneous mechanisms is stochastic resonance, in which cells whose membrane potential is near firing threshold can be induced to fire by the addition of an external field (Ozen et al., 2010). Related to stochastic resonance, the “rhythm resonance” mechanism involves precisely timed fields that are coordinated with the depolarizing phase of neurons whose membrane potential is spontaneously fluctuating in an oscillating fashion (Deans et al., 2007; Frohlich and McCormick, 2010; Reato et al., 2010, 2013). That is, tACS may act by summation with the membrane potential of active neurons such that when tACS is in-phase with the ongoing or membrane potential fluctuations, spiking rates increase. External fields may combine with internal fields in such a way to cluster spiking, without actually increasing or decreasing the overall firing rate (Liu et al., 2018). Finally, external fields of high strength can entrain local networks by overwhelming the ongoing activity (Liu et al., 2018). Proposed non-instantaneous mechanisms include various forms of synaptic plasticity that effect synaptic efficacy (Stagg and Nitsche, 2011; Stagg et al., 2018).

Of present interest, tACS has been paired with periodic sensory stimulation that is expected to drive cortex at a specific frequency and its harmonics. This approach has the potential to provide strong evidence for instantaneous or “on-line” effects, as well as non-instantaneous (“off-line”) effects. An example of the former is the finding that hearing threshold for periodic tone bursts can be modulated by frequency-matched tACS in a phase-dependent fashion (Neuling et al., 2012).

Here we used tACS and periodic visual stimulation to measure online and offline effect of tACS on visual perception, using the phenomenon of binocular rivalry as a perceptual read-out. Binocular rivalry occurs when images presented to each eye are not fusible into a single percept. Instead of seeing a single image, as in normal viewing, perception alternates between the images presented to each eye. We reasoned that if tACS modulates neural activity in a frequency or phase-specific fashion, it could alter the balance of perceptual salience for images presented to each eye and this alteration may be reflected in the statistics of the perceptual alternations. Importantly for present purposes, the dynamics of the perceptual alternation depend on the relative perceptual strength of the two images and these dynamics have been formalized into a set of laws which can be used as an interpretive framework (LeVeldt, 1965; Brascamp et al., 2015). We test both the phase and frequency specific models of tACS, looking to see if tACS might have constructive or destructive effects when presented in phase or out of phase with visual stimulation of the same frequency, or if merely being frequency matched is all that is required of tACS for it to have an effect, either online or offline.

## METHODS

### Participants

Participants in this study were 34 females and 35 males from Stanford and the surrounding community who received either credit in a psychology course or monetary payment of $30. Participants were screened to have a stereo acuity better than 50 arc-seconds on the RanDot stereo-acuity test, averaging 31.28 arc-seconds. Participants had an average right eye visual acuity of −0.068 and an average left eye visual acuity of −0.061 on the Bailey-Lovie chart. Eye dominance was measured with the hole in the card test and revealed that 19 participants were left eye dominant and 50 were right eye dominant. Procedures were approved by the Stanford Institutional Review Board and each participant provided written informed consent.

### Visual Stimulation

For each of three experiments, 10% Michelson contrast, dichoptic Gabor patches (3 c/deg, sigma 1 deg) were presented to the participants. The Gabor patches were oriented at 135 degrees in the left eye and 45 degrees in the right eye. The patches flashed either at 3Hz or 5Hz, depending on the experiment. The stimulus was generated using in-house software and presented on a 55-inch, LG passive 3D OLED screen at a mean luminance of 10cd/m^2^. The Gabor patches were viewed by participants from 80cm from the stimulus display. Three concentric circles, visible to both eyes acting as a fusion and focus lock surrounded the test stimulus. Participants wore polarizing-filter glasses and placed their head on a head-rest that had been pre-measured to be 80cm from the stimulus display. The experiments were performed in a darkened room and participants signaled their rivalry percepts by via button presses. They were instructed to press the left button if the bars on the screen appeared to be oriented up and to the left, and the right button if the bars on the screen appeared to be oriented up and to the right. They were instructed to press neither button in the case of mixed percepts. They were also given a practice period and the opportunity to ask for any clarification.

In the first two experiments described below, the stimulus flashed at 3Hz in each eye, with left eye stimulus being shifted in phase by 180 degrees relative to the right eye. In the third experiment, the left eye visual stimulus was presented at 5Hz and the right eye stimulus was presented at 3Hz. Electrical stimulation, when present was delivered at 3 Hz.

### tACS Stimulation

Sinusoidal electrical stimulation was delivered via a StarStim system (NeuroElectrics, Barcelona). For every participant, the protocol before starting stimulation involved: hair-washing with baby shampoo, electrode cap sizing, abrasive skin preparation via Lectron II gel application to clean the targeted scalp regions, electrode placement, and electrode gel application. Impedances were kept below 10k. The two source electrodes were placed at P07 and P08, while the common sink electrode was placed at O_Z_. Stimulation current amplitudes were 1 mA at each of the source electrodes and 2 mA at the sink electrode. This “tri-polar” montage was designed to minimize current flow through the eyes and thus to minimize the chance of inducing phosphenes (Kar and Krekelberg, 2012) and to focus the stimulation on the central field (∼2 deg) representations in V1, V2 and V3 that are located on the occipital pole(Dougherty et al., 2003). Current distributions were visualized with StarStim BEM modeling software.

The StarStim system was controlled over Bluetooth. Because the start time of the electrical stimulation upon initiating a request for stimulation over Bluetooth was not deterministic, the computer generating the visual stimulus monitored the phase of the tACS stimulation in real time by recording the differential signal between electrodes at Oz and PO8. The visual-stimulation software monitored the tACS waveform in real time and scheduled the start of visual-stimulation so it began at each of 4 pre-determined phase offsets of 0, 90, 180 or 270 degrees relative to the tACS stimulus. This allowed the system to match the current and visual stimulus phase relationships with a maximal error of 1 video frame duration (16.6 msec or 18 deg of phase at 3 Hz).

Electrodes were also monitored via the Bluetooth stimulator, which displayed real-time feedback on the impedance and waveforms of the stimulating current, so that the experimenter could visualize the functionality and impedance of all three electrodes and the output of the tACS stimulator that was connected to the electrodes.

### Experimental procedure

The protocols were designed to assess rivalry before, during and after tACS, testing whether tACS is phase or frequency specific. We also included a control experiment to determine the stability of rivalry parameters over the duration of the active tACS experiments. This psychophysical control was used to determine whether there was any “drift” in the rivalry process after being exposed to the same stimulus for the duration of the experiment. In this control experiment, referred to below as the no-tACS control, the stimulus conditions and experiment duration were identical to the ones to be described for the first tACS experiment (3 Hz anti-phase visual stimuli), but no electrical stimulation was applied at any time.

A pilot study was conducted to investigate whether orientation (45/135 deg), temporal frequency (3Hz/5Hz) or eye (left/right) biased overall rivalry distributions. We measured rivalry durations for each of the 8 pairwise combinations in 5 observers. By testing these various iterations of the same paradigm, we established that there was no effect of any of the three factors on overall rivalry modulation, allowing us to pick fixed pairings of eye and orientation and a fixed targeted eye.

In each experiment, an initial, non-stimulated block was presented to measure the participant’s baseline rate of rivalry. This first block lasted 4 minutes and was comprised of two trials, each lasting approximately two minutes. Next, blocks with 4 different tACS relative phases were run consecutively, but in random order for every subject to control for temporal order. Each block contained 4 trials, lasting approximately two minutes. For the duration of these blocks, participants received tACS stimulation at four different relative phases: 0°, 90°, 180°, or 270°. This portion of the experiment lasted 28 minutes. tACS stimulation was on for the entire 28-minute period. Immediately after the completion of the four blocks with tACS stimulation, participants completed a 6^th^, non-stimulated block of two trials, lasting 4 minutes. Participants were then given a 10-minute break in which they could wash their hair but not leave the room. After the 10 minutes, participants completed condition 7, which also had no stimulation but featured the exact same stimulus. This was done in order to measure a possible persistent effect of tACS.

In the first tACS experiment (Experiment 2) and in the psychophysical control experiment, both Gabor patches flashed at 3Hz, but they were offset by 180° of one another. This made the sign of polarization opposite in the two eyes and was our test of phase specificity. During blocks 2-5, tACS was presented at 3Hz for the entire duration. When electrical stimulation was applied, tACS was synchronized with the 3Hz visual stimulus, such that it was at either 0°, 90°, 180°, or 270° offset in relation to the visual stimulus in the right eye.

In Experiment 3, the visual stimulus was presented at two different frequencies. This was our test of frequency specificity. The right eye’s Gabor patch flashed at 3Hz, while the left eye’s Gabor patch flashed at 5Hz. tACS was presented at 3Hz, targeting the right eye. During blocks 2-5, when tACS stimulation was applied, the tACS was synchronized with the 3Hz stimulus, such that it was at 0°, 90°, 180°, or 270° phase in relation to the visual stimulus. The relative phase of tACS with respect to the right eye precessed through all possible phase angles.

### Data Preprocessing

The raw data was exported into MATLAB files for analysis. After exporting, data was then sorted into left and right presses. Next, duration lengths were calculated for each subject and each trial. The button hardware occasionally resulted in very short durations of non-press during periods of button press. These durations were cleaned up by consolidating any user non-press that lasted <0.5 seconds to the previous button press, provided it was the same button. Figure 2 shows raw button durations for three observers each with very different duration distributions spanning the range observed in the experiment. To characterize these distributions, all data was transformed into log-seconds and then fitted utilizing a normal distribution. The log-normal fits were better than traditional gamma fits.

**Figure 1:**
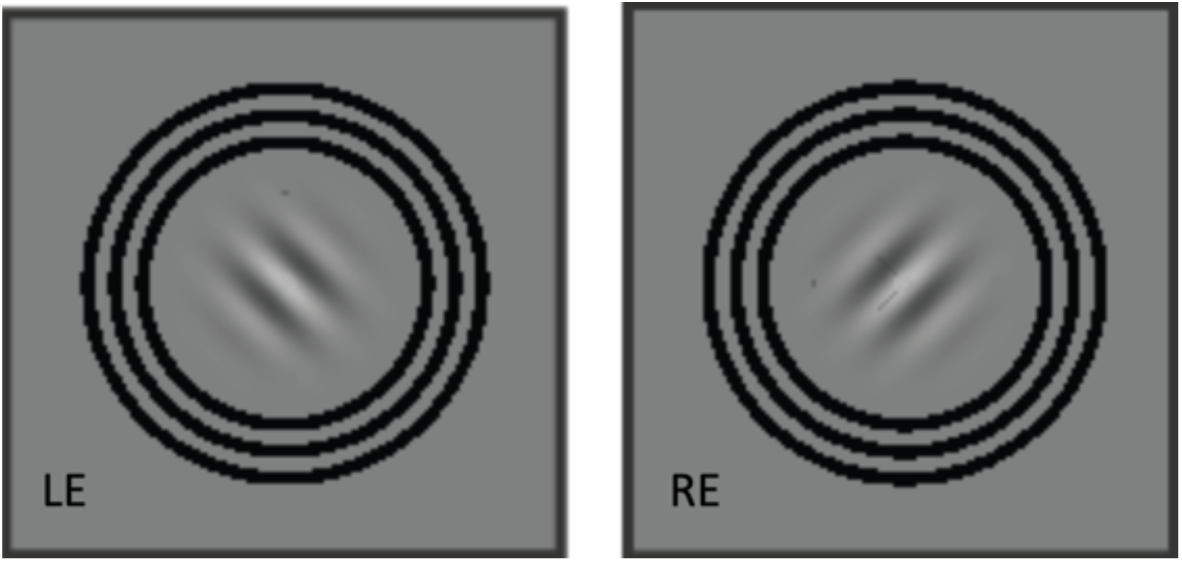
Visual stimulus schematic. The two Gabor patches with orthogonal carrier orientations were overlaid on one another on the 3D screen. The left eye saw the image on the left, while the right eye only saw the image on the right.

**Figure 2:**
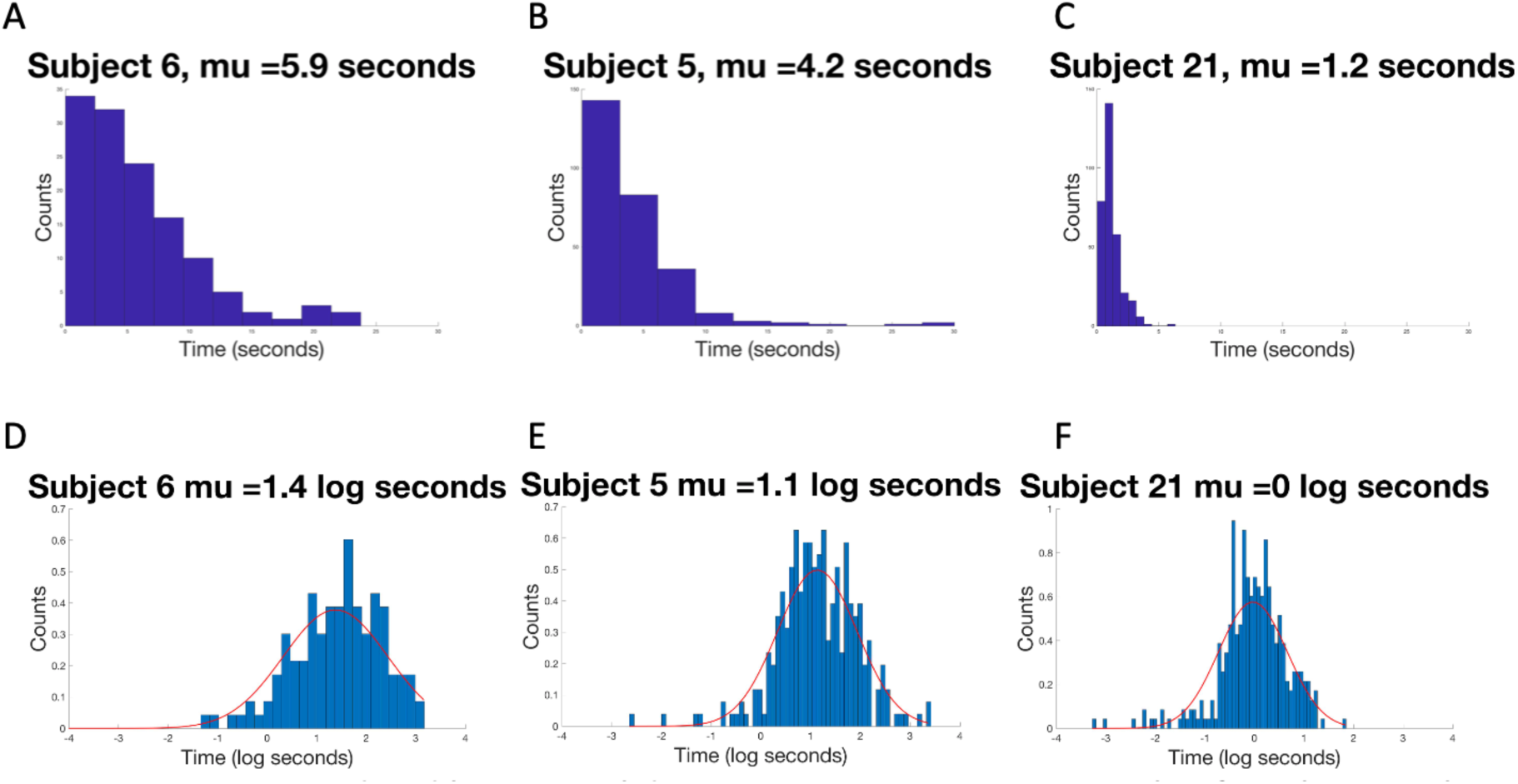
Unprocessed and log-normalized data. A: Representative example of a participant with a longer mean rivalry duration. B: Representative example of a participant with an average mean rivalry duration. C: Representative example of a participant with a shorter mean rivalry duration. D-F: Log-transforms of the corresponding distributions A-C, demonstrating that the log-normal fit (red curve) accurately represents data for all rates of rivalry.

### Outlier exclusion

We noted that some participants had extremely long rivalry durations, suggesting that their rivalry process was far from being balanced. As this imbalance would likely reduce the sensitivity of the rivalry read-out to tACS perturbation, we developed an outlier screening procedure based on the raw durations. A 1.5IQR test was utilized to exclude outliers from each of the three experiments. For this exclusion, only means from condition 1 in each experiments were considered. This was done to ensure that we were excluding subjects who didn’t have typical rivalry alternations in their baseline reading, not running the risk of excluding someone who might have been significantly affected by the electrical stimulation.

### Data normalization

As can be seen from Figure 2, the mean of each subject’s log-durations varied significantly significant variation between the baseline mean durations. This variation arose because of the variable rate at which the subjects rivaled, with some rivaling faster than others. As the goal of the experiment is to assess the effects of tACS on perception within participants, we normalized all means to the participants value in the baseline condition by subtracting each condition’s values from that of condition 1. Normalization was performed for both the log mean duration and switch-rate metrics. The effect of the normalization on duration measurements is shown in Figure 4 which plots the data from all conditions for the three observers shown in Figure 2. Panels A-C show the un-normalized log durations that have a larger range than after normalization (panels D-F).

**Figure 3:**
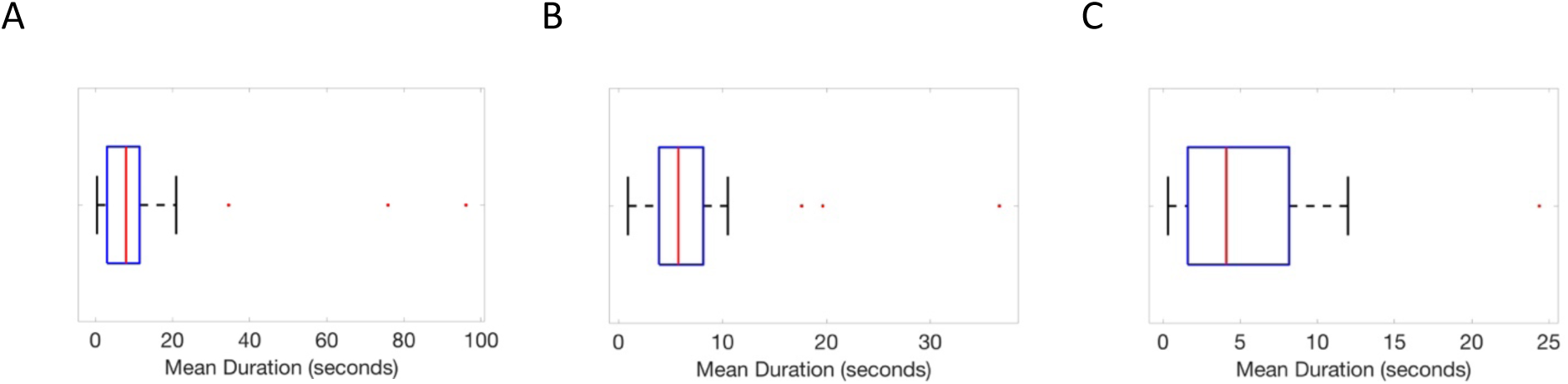
Outlier detection. Detection of mean durations that fell outside of the pre-determined 1.5IQR of the data. A: Box and whisker plot for the 3Hz/5Hz visual stimulation Experiment. B: Box and whisker plot for the 3 Hz anti-phase visual stimuli setup. C: Box and whisker plot for the no-tACS control experiment. Outliers were data points that fell beyond 1.5*IQR + Q3 and are indicated by the red dots outside of the boxplot. These participants (n=3, 3 and 1 for A, B and C, respectively) were excluded from further analysis, as they did not show typical rivalry alternations.

**Figure 4:**
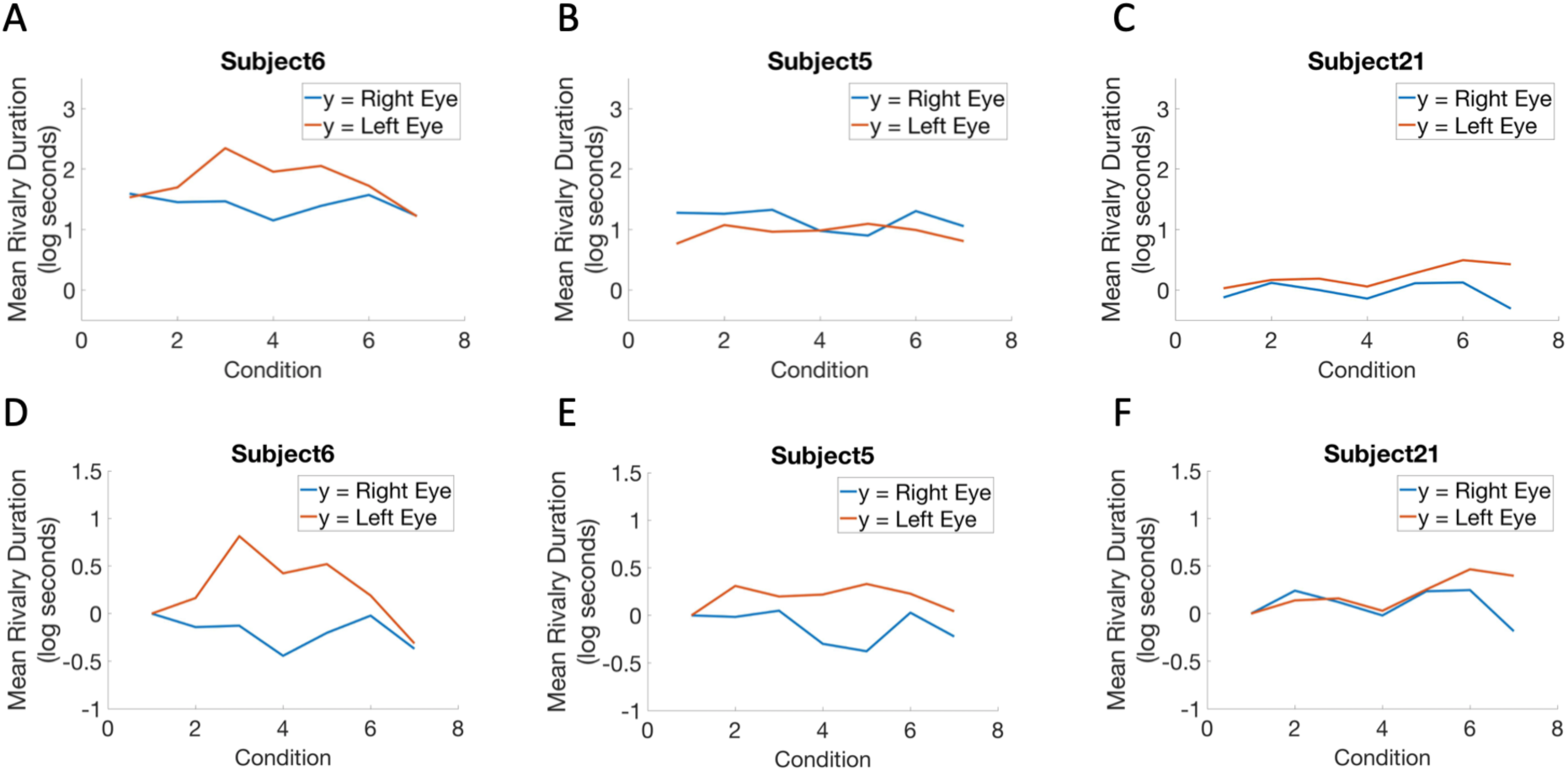
Normalizing duration data to a set baseline. A-C: Representative participants (same as those shown in Figure 2) showing the variability in the baseline and overall rivalry durations. D-F: Normalized participant data obtained by setting condition 1 as the subtracted baseline.

Once normalized, these data were averaged to generate graphs of mean normalized duration for right and left button presses and graphed over condition. For each condition, we also calculated the rivalry switch rate by counting how many times each subject switched buttons throughout the experiment and dividing that number by the duration of the experiment.

### Statistical Analysis

To assess the impact of the phase of tACS on rivalry, paired t-tests were performed for all pair-wise combinations of the four test conditions, and then corrected utilizing the Bonferroni adjustment for multiple comparisons. If these tests revealed no significant differences, data from the four test conditions were pooled and tested against zero. This test determined the effectiveness of tACS at modulating rivalry in a phase-independent manner.

For the post-tACS conditions, paired t-tests compared condition 6 to condition 7 (immediately post-tACS to 10 minutes post-tACS) to determine if there were any significant lasting effects of the stimulation. The two conditions were also t-tested against zero (the baseline condition) to see if there was any overall effect of tACS. If the t-tests revealed that conditions 6 and 7 were statistically indistinguishable, the two conditions were pooled first and then compared to zero.

## RESULTS

### Behavioral control experiment without tACS

This experiment was performed to determine the degree to which rivalry features remain constant over the long duration of the tACS experiments. Drift in rivalry parameters over the duration of the experiment, if it occurred, would complicate the interpretation of the tACS data. Therefore, to test rivalry stability, we used the same number of trials and experiment duration as in the tACS experiments. The participants viewed 3 Hz anti-phase stimuli in each eye and reported which eye was dominant by holding down one button for left eye and a second button for right eye dominant periods.

We observed no significant deviations from baseline durations or switch rates over the course of the experiment which lasted ∼50 minutes. Note that conditions 2-5 were presented in random over a period of 28 minutes, so to determine whether rivalry durations change over the period used in the active tACS phases of Experiments 2 and 3, we pooled the data across conditions 2-5 to obtain estimates of duration and switch rates at an average of 14 minutes after baseline. The immediate post-test (condition 6 data reflect rivalry at and average of 30 minutes after baseline and the final post-test (condition 7), 42 minutes. In each case we tested the durations against the baseline value of zero. Each of the pooled condition 2-5 duration values were non-significant (p=0.33; left eye, p= 0.55. right eye). Similarly tests of the periods corresponding to the post-tACS conditions also revealed no significant changes in mean duratione from baseline. For the immediate post-test the right eye p-value was 0.95 and the left eye vlaue was 0.88. The corresponding values for the 10-minute post-tACS condition were 0.96 and 0.77, respectively). For the switch rate parameter estimated at 14 minute, there was no measurable deviation from baseline for either eye (p=0.83), nor were there measurable changes in the immediate post-test (p=0.70) or final post-test (p=0.39).

### Phase-specificity of tACS

The purpose of the next experiment was to determine whether tACS can modulate rivalry durations or switch rates in a phase-specific fashion. To make this measurement, we presented 3Hz visual stimulation to both eyes, but with 3Hz anti-phase tACS. Because we don’t know the precise timing of the visual activity we are reading out, we presented tACS at 4 different relative phases to span one full cycle of the visual and tACS stimulation (blocks 2-5). In each case tACS was in antiphase, given that the visual stimuli were always presented in anti-phase to the two eyes. The phase-specific hypothesis predicts that a tACS phase that is maximally suppressive in one eye will be maximally facilitative in the other eye which is receiving the opposite polarization, and vice versa, once the proper absolute delay between the visual system’s internal response and the instantaneous tACS field is matched. We defined the relative phase of tACS with respect to the right eye’s visual stimulation. If tACS was phase specific, we should see weakening in one eye accompanied with a strengthening in the other eye at a particular absolute phase value and the opposite effect at a phase value shifted by 180 deg.

The data for this experiment are presented in Figure 6, with left-eye data plotted in red and right-eye data plotted in blue for each block of the experiment. We first tested whether relative tACS/visual phase affected rivalry durations by performing all pair-wise comparisons between the four relative phase values (conditions 2-5 in Figure 6). We observed no phase-specific effect during the application of tACS. All Bonferroni adjusted p-values for the left and right eye pair-wise comparisons were 1.

**Figure 5:**
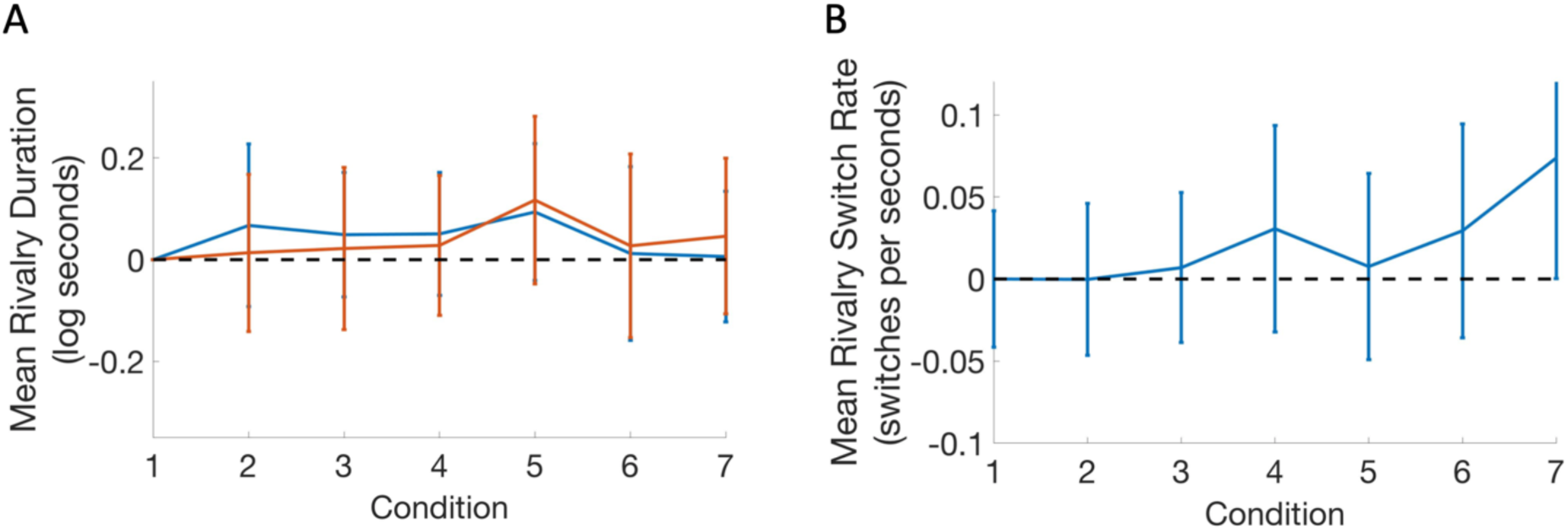
No tACS control experiment: 3 Hz right eye stimulation, 3Hz left eye stimulation without tACS stimulation. Panel A plots left-(red) and right-eye (blue) normalized mean durations. Panel B plots normalized mean rivalry switch rates over blocks of the experiment. Error bars plot +/- 1 s.e.m.

**Figure 6:**
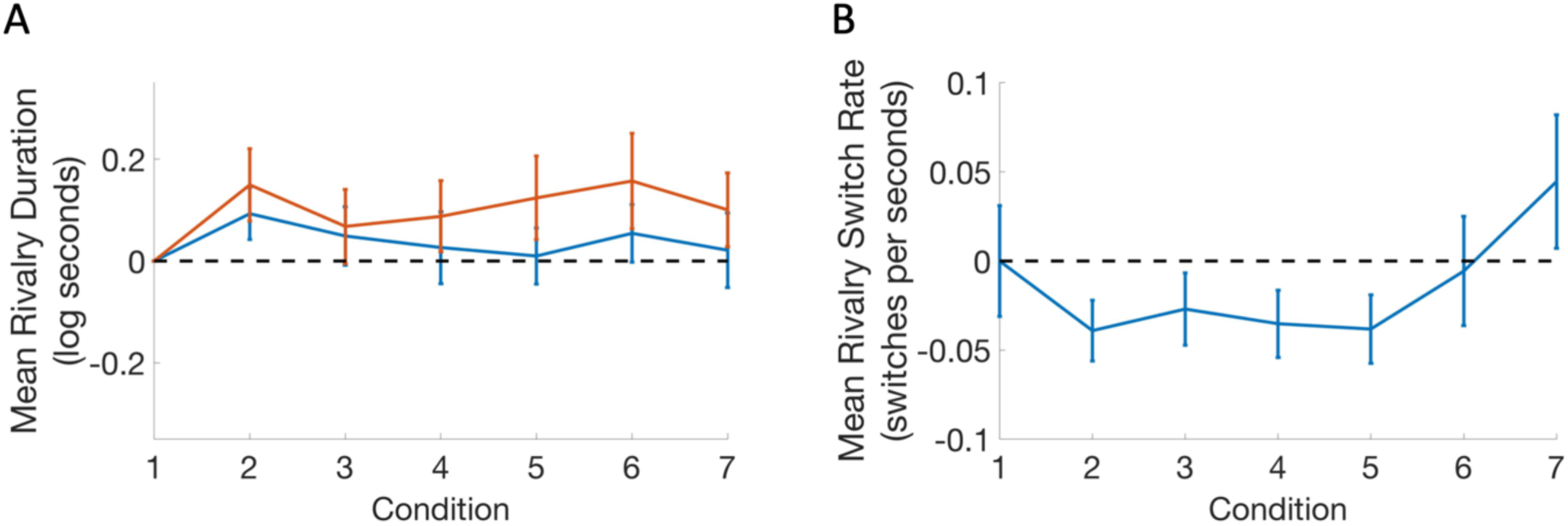
3 Hz right eye stimulation, 3Hz left eye stimulation with 3Hz anti-phase tACS. Panel A plots left-(red) and right-eye (blue) mean durations B: Mean rivalry switch rate over blocks of the experiment. Error bars plot +/- 1 s.e.m.

While tACS did not modulate rivalry in a phase-dependent fashion, it could still exert an effect that is independent of relative phase. To test for this possibility, we pooled all of the mean duration measurements for each eye made during tACS stimulation and compared them to 0. The mean durations increased by a factor of 1.28 in the left eye compared to baseline and by a factor of 1.11 in the right eye. The shift for the left eye was highly significant (p= 0.004) but was marginally significant for the right eye (p= 0.13).

Another important parameter of the rivalry distribution is the rate of switches. Figure 6B shows the rivalry switch rates. All pair-wise Bonferroni corrected tests over the tACS-present conditions had p-values = 1 and thus showed no phase specificity. By contrast, switch rates during tACS decreased by a factor of 1.22 (p< 0.000).

We then asked whether any of the effects on mean duration and switch rate observed during tACS persisted immediately after cessation of tACS or after 10 minutes after tACS cessation. Mean durations for the immediate post-tACS condition were not significantly different from baseline for the right eye (p=0.35), but there was a non-significant trend for longer durations in the left eye (p=0.11). There were no measurable effects on mean duration for the 10-minute post-tACS condition (right eye 0.78, left eye 0.18). Switch rates in the immediate post-tACS condition were not measurably different from baseline (p=0.90), nor were they different from baseline in the 10-minute post-tACS condition (p=0.36).

### Frequency Specificity of tACS

Having found no phase specificity of tACS stimulation, we moved on to test whether the tACS effects were frequency specific. This experiment visually stimulated the right eye at 3Hz and the left eye at 5Hz. We applied 3Hz tACS during the four middle “test” conditions and again varied tACS/visual phase. Thus, the right eye received synchronous 3 Hz tACS stimulation at different relative phases, but the left eye received asynchronous tACS/visual stimulation as the visual stimulus was presented at 5 Hz. If tACS’ effects were frequency specific, the effects of tACS should be most apparent in the synchronously stimulated right eye.

The data from this experiment are shown in Figure 7. As in Experiment 2, we observed no phase-specific effects on mean rivalry duration during the application of tACS. All pair-wise, Bonferroni-corrected p-values were 1 for both eyes. We thus pooled all of the mean duration measurements for each eye made during tACS stimulation and compared them to 0 to detect any significant modulation from baseline in the two eyes’ percepts. The mean durations increased by a factor of 1.08 in the left eye and *decreased* by a factor of 1.50 in the right eye. The shift for the right eye was highly significant (p< 0.000) but was marginally significant for the left eye (p= 0.19). Figure 7B shows the rivalry switch rates. The switch rates did not differ by phase on pair-wise Bonferroni corrected tests (all p-values = 1), but they decreased overall by a factor of 1.28 (p< 0.000). The fact that the two eye durations change in opposite directions in this experiment, versus changing in the same direction as 3 Hz anti-phase experiment, plus the fact that switch rates decrease suggests tACS is effective and is frequency specific.

**Figure 7:**
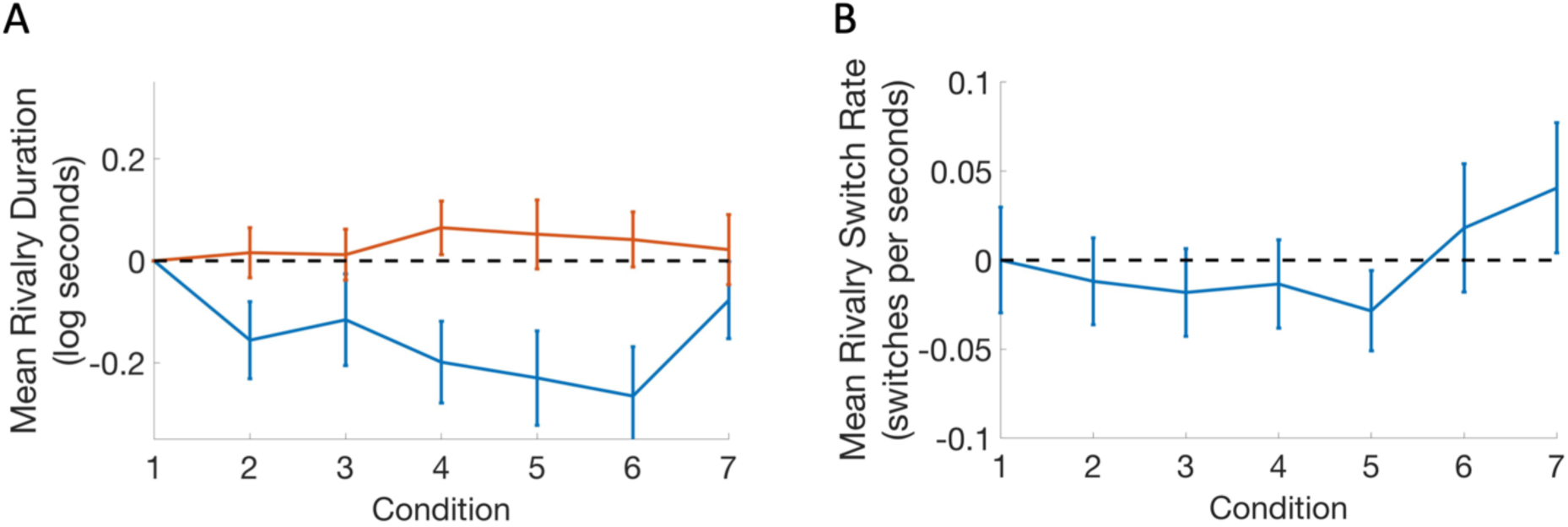
3 Hz right eye visual stimulation, 5Hz left eye visual stimulation. 3Hz tACS reveals a slowing of percept durations in targeted eye and no change in nontargeted eye. Panel A shows rivalry durations for both left (orange) and right (blue, targeted) eyes. Panel B shows rivalry switch rates. Rivalry slows in the targeted eye, and switch rates decrease.

We then asked whether any of the effects on mean duration and switch rate observed during tACS persisted immediately after cessation of tACS or after 10 minutes post tACS. Mean durations for the immediate post-tACS condition were significantly different from baseline for the right eye (p=0.01) but were not significantly different from baseline for longer durations in the left eye (p=0.44). There were no measurable effects on mean duration for the 10 minute post-tACS condition (right eye 0.32, left eye 0.75). Switch rates in the immediate post-tACS condition were not measurably different from baseline (p=0.78), nor were they different from baseline in the 10-minute post-tACS condition (p=0.45).

## DISCUSSION

Our results using bi-stable rivalry percepts as a readout suggest that the effects of tACS on visual perception depend on matching the frequency of visual and tACS stimulation, but not on the relative phase of matched-frequency stimulation. The lack of phase specificity was found in both the 3 Hz anti-phase and 3Hz/5Hz experiments. When 3Hz/5Hz frequencies of visual stimulation were used, tACS that matched the visual stimulation rate (3Hz) decreased the mean durations in that eye, and there was a marginally significant increase in duration in the other (5Hz). By contrast, when anti-phase tACS of the same frequency (3Hz) was presented to each eye, both eyes durations increased. Together, these results indicate that the effect of tACS was frequency, but not phase specific. Levelt’s laws of rivalry allow us to deduce that that tACS acted to reduce internal response strength, as described in the next section.

Our findings are largely interpretable within the framework of Levelt’s laws governing rivalry as recently updated by Brascamp and co-workers (LeVeldt, 1965; Brascamp et al., 2015). Levelt’s original laws were based on the results of psychophysical studies of rivalry including situations that varied the relative stimulus intensities in the two eyes. Here, we consider whether tACS effectively modulates internal response strength in a way similar to changes stimulus intensity. There are four Levelt’s laws and we interpret our data with respect to each of them in the order that they have been traditionally stated. Levelt’s first law states that increasing the stimulus strength in one eye increases the perceptual dominance of that eye’s stimulus, and conversely, decreasing the stimulus strength in one eye decreases its perceptual dominance. Here we do not manipulate visual stimulus strength, but we sought to manipulate the strength of the internal neural response via tACS. In our test of frequency-specificity with 3Hz/5Hz stimulation, we observed a phase-independent decrease in the right eye’s perceptual dominance reflected by its shorter mean durations consistent with this eye’s internal representation being weakened. Weakening of the representation in traditional rivalry experiments would come about by decreasing stimulus intensity, so we interpret the effect of tACS in this experiment as being effectively suppressive.

Levelt’s second law states that “increasing the difference in stimulus strength between the two eyes will primarily act to increase the average perceptual dominance duration of the eye with the stronger stimulus.” In the 3Hz/5Hz experiment, tACS targeted the right eye, acting to shorten its durations, consistent with its response being weakened based on the above prediction from the first law. According to the second law, this should have the effect of increasing the mean duration of the opposite eye. We see a trend in this direction, but the effect is larger on the targeted eye. While the original second law stipulated that this increase in the non-targeted eye should have been larger than the effect on the (weakened) targeted right eye, more recent findings have shown that the second law is only valid for a relatively restricted range of stimulus intensities and thus internal response intensities, as studied here and can even reverse sign (Brascamp et al., 2006). Second law predictions are thus not strongly proscriptive in the present context.

Levelt’s revised third law states that “increasing the difference in stimulus strength between the two eyes will reduce the perceptual alternation rate” (Brascamp et al., 2015). The data from 3Hz/Hz experiment are relevant here and we indeed see a reduction in the perceptual alternation rate from baseline when the two eyes are stimulated asymmetrically, *e.g*. only the right eye received synchronous tACS stimulation. Related to this, bistable percept alternation rates have been shown to be maximal when competing percepts are perceptually balanced, as they would have been at baseline (Moreno-Bote et al., 2010). A modulatory effect of tACS (in either direction) is thus predicted to lower the alternation rate, which is what we observe.

Levelt’s fourth law states that “increasing stimulus strength in both eyes while keeping it equal between eyes will generally increase the perceptual alternation rate….” (Brascamp et al., 2015). The converse is true and more relevant to our observations. Switch rates decrease in the 3Hz/3Hz condition, suggesting that internal response strength decreased, which is consistent with mean durations having also decreased, as suggested above for the first law interpretation of the mean duration data of the 3Hz/5Hz experiment which used asymmetric stimulation.

### Possible mechanisms underlying observed tACS effects

Prior work in animal models has suggested that tACS can exert phase-specific effects via interactions between the imposed field and cell membrane potentials (Deans et al., 2007; Frohlich and McCormick, 2010; Reato et al., 2010, 2013). Two non-invasive studies in human have varied the relative phase of synchronous tACS and periodic sensory stimuli. The first study (Neuling et al., 2012) reported that psychophysical thresholds for detecting brief 500 Hz tones in noise depended on the timing of the tones relative to 10 Hz tACS. This effect was subsequently replicated with 4 Hz tACS (Riecke et al., 2015). A similar approach has not been taken in the visual modality, to our knowledge. In the visual modality, Ruhnau and co-workers (Ruhnau et al., 2016), presented visual flicker at 7 and 11 Hz in presence of frequency-matched or non-matching tACS. They found tACS effects to be frequency-specific, as we report here. They could not assess phase-specificity as their visual and electrical stimuli were not phase-synchronized as they were in the present experiments.

The relative lack of phase-specificity in our experiment relative to that reported previously with auditory stimuli could arise for a number of different reasons. First, the effective current strength delivered to cortex could differ for auditory vs visual stimulation montages. The auditory experiments used a bipolar montage with electrodes over auditory cortex where the skull is much thinner than it is over the occipital pole where our 3-electrode array was located. Instantaneous, phase-specific modulation effects are expected to depend strongly on delivered field strength and recent estimates have suggested that the deposited field in human transcranial stimulation experiments is weak relative to that needed to influence membrane potential (Opitz et al., 2016; Huang et al., 2017; Voroslakos et al., 2018). Secondly, the rivalry stimuli were highly supra-threshold, rather than being near threshold as in the auditory detection experiments. tACS effects may depend on the strength of the internal response, which is very small for near threshold stimuli and larger for supra-threshold stimuli. Network-level activity may thus have differed substantially. Finally, the effect of tACS not only depends on electrical path length between the electrodes and the active tissue, but it also depends on tissue orientation (Opitz et al., 2011). We attempted to minimize this effect by using small visual stimuli that will produce relatively small patches of cortical activation. Because there are multiple representations of the visual field, the separate representations of the patches in different visual areas may have sufficiently different orientation to eliminate the effect of tACS phase which could invert depending on tissue orientation.

Transcranial Electrical Stimulation can also act via non-instantaneously through mechanisms that modulate synaptic efficacy (Zaehle et al., 2010; Jackson et al., 2016; Kronberg et al., 2017). These effects are necessary to explain the offline effect we have observed, as there is no imposed field present to interact with cell membrane potentials in the offline conditions. Recall that we showed an immediate offline effect on targeted-eye rivalry duration in the 3Hz/5 Hz condition. It is possible that these mechanisms also underlie the effects we observe during tACS stimulation. Consistent with this, the sign of the offline effect is the same as the effect during tACS stimulation.

In conclusion, tACS stimulation can modulate visual perception and under our conditions its net effect is consistent with a weakening of the internal visually-driven signal.

## Acknowledgements

Supported by 1R21EY026748 from the National Eye Institute, National Institutes of Health, Stanford Bio-X Summer Undergraduate Research Program.

